# A major locus controls local adaptation and adaptive life history variation in a perennial plant

**DOI:** 10.1101/178921

**Authors:** Jing Wang, Jihua Ding, Biyue Tan, Kathryn M. Robinson, Ingrid H. Michelson, Anna Johansson, Björn Nystedt, Douglas G. Scofield, Ove Nilsson, Stefan Jansson, Nathaniel R. Street, Pär K. Ingvarsson

## Abstract

**Background:** The initiation of growth cessation and dormancy represent critical life-history tradeoffs between survival and growth, and have important fitness effects in perennial plants. Such adaptive life history traits often show strong local adaptation along environmental gradients but despite their importance, the genetic architecture of these traits remains poorly understood.

**Results:** We integrate whole genome re-sequencing with environmental and phenotypic data from common garden experiments to investigate the genomic basis of local adaptation across a latitudinal gradient in European aspen (*Populus tremula*). We discover a single genomic region containing the *PtFT2* gene that mediates local adaptation in the timing of bud set and that explains 65% of the observed genetic variation in bud set. This locus is the likely target of a recent selective sweep that originated right before or during colonization of northern Scandinavia following the last glaciation. Field and greenhouse experiments confirm that variation in *PtFT2* gene expression affect the phenotypic variation in bud set that we observe in wild natural populations.

**Conclusions:** Our results reveal a major effect locus that determine the timing of bud set and that have facilitated rapid adaptation to shorter growing seasons and colder climates in European aspen. The discovery of a single locus explaining a substantial fraction of the variation in a key life history trait is remarkable given that such traits are generally considered to be highly polygenic. These findings provide a dramatic illustration of how loci of large-effect for adaptive traits can arise and be maintained over large geographical scales in natural populations.

## Backgrounds

Most species are distributed over heterogeneous environments across their geographic range and spatially varying selection is known to induce adaptation to local environments [1]. Local adaptation thus provides an opportunity to study population genetic divergence in action [2]. Although the interaction between gene flow and natural selection is well studied from a theoretical point of view and makes a number of testable predictions [3], there are to date few empirical studies investigating how local adaptation is established and maintained at the molecular level in natural populations.

Many perennial plants, such as forest trees, have wide geographic distributions and are consequently exposed to a broad range of environmental conditions, making adaptation to diverse environmental and climate conditions crucial in these species [4-7]. Natural populations of these plants are often locally adapted and display pronounced geographic clines in phenotypic traits related to climatic adaptation even in the face of substantial gene flow [5,6]. One of the most important traits mediating local adaptation is initiation of growth cessation at the end of the growing season, which represents a critical life history trade-off between survival and growth in most perennial plants [8,9]. Local adaptation in phenology traits, such as growth cessation, is well documented at the phenotypic level in many long-lived perennial species [2,6]. Compared to traditional model and crop species that are usually annuals, naturally inbred and have rich genomic resources available, the genomic and evolutionary research in long-lived, outcrossing perennial species are much more difficult to conduct, and the genetic architecture of adaptive traits in such species is therefore still rather poorly understood [5,6].

Here we investigate the genomic signatures of local adaptation across a latitudinal gradient determining the length of the growing season in European aspen (*Populus tremula*). *P. tremula* is a dioecious and obligately outbreeding tree species, and both seeds and pollen are wind-dispersed, resulting in frequent long-distance gene flow and consequently weak population structure [10,11]. Despite extensive gene flow, local populations display strong adaptive differentiation in phenology traits, such as the timing of bud set and growth cessation, across the latitudinal gradient[10]. In this study, we integrate whole genome re-sequencing with field and greenhouse experiments to characterize the genome-wide architecture of local adaptation in *P.tremula*. Using a combination of approaches we identify a single genomic region, centered on a *P. tremula* homolog of *FLOWERING LOCUS T2* (*PtFT2*), that controls a substantial fraction of the naturally occurring genetic variation in the timing of bud set. The region displays multiple signs of a recent selective sweep that appears to have been restricted to the northernmost populations. Our results provide evidence of a major locus that has facilitated rapid adaptation to shorter growing seasons and colder climates following post-glacial colonization.

## Results

### Genome sequencing, polymorphism detection and population structure

In this study we used a total of 94 unrelated *P. tremula* trees that were originally collected from twelve sites spanning c. 10 degrees of latitude (~56-66°N) across Sweden (the SwAsp collection from [12], see also Additional file1: Table S1). Earlier studies have shown that the SwAsp collection display a strong latitudinal cline in the timing of bud set (Fig. 1a,b) [10-12]. We performed whole genome re-sequencing of all 94 aspens and obtained a total of 1139.2 Gb of sequence, with an average sequencing depth of ~30 × per individual covering more than 88% of the reference genome (Additional file1: Table S1). After stringent variant calling and filtering, we identified a total of 4,425,109 high-quality single nucleotide polymorphisms (SNPs) with a minor allele frequency (MAF) greater than 5%.

**Figure 1.**
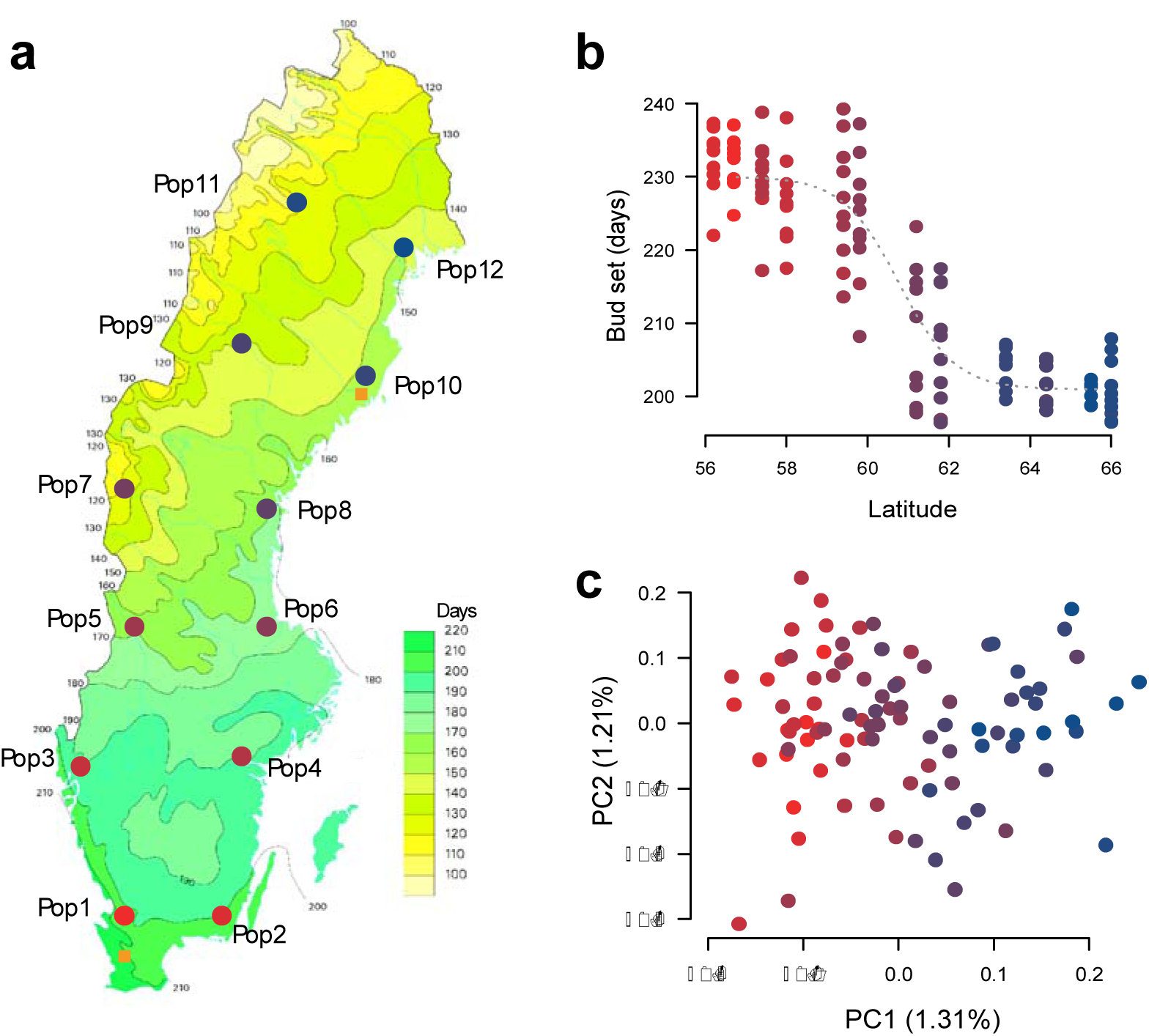
Geographic distribution and genetic structure of 94 aspen individuals. a) Location of the twelve original sample sites of the SwAsp collection (circles) and the location of the two common garden sites (squares). The original collection sites span a latitudinal gradient of c. 10 latitude degrees across Sweden. b) Breeding values for date of bud set for the 94 individuals included in the study across the two common gardens and three years (2005, 2006 and 2007). c) Population structure in the SwAsp collection based a principle component analysis of 217,489 SNPs that were pruned to remove SNPs in high linkage disequilibrium (SNPs included all have *r*^2^<0.2). Although two axes are shown, only the first axis is significant (*P*=3.65×10^−12^, Tacey-Widom test, 1.31% variance explained).

We found very weak population structure across the entire range using principal component analysis (PCA) [13], with a single significant axis separating individuals according to latitude (*r*=0.889, *P*-value <0.001) but explaining only 1.3% of the total genetic variance (Fig. 1c; Additional file2: Table S2). Consistent with this, a Mantel test also showed a weak pattern of isolation by distance (*r*=0.210; *P*-value=0.047; Additional file3: Figure S1). Swedish populations of *P. tremula* have gone through a recent admixture of divergent postglacial lineages following the Last Glacial Maximum (LGM) [14] and it is possible that this is capable of generating a genome-wide pattern of clinal variation. However, the extremely low genetic differentiation we observe among populations (mean *F_ST_*=0.0021; Additional file3: Figure S2) suggests that extensive gene flow within *P. tremula* has almost eradicated any such signal across the genome.

### Identifying genomic variants associated with local adaptation

We used three complementary approaches to identify candidate SNPs involved in local adaptation. First, we identified SNPs that were most strongly associated with the observed population structure using PCAdapt [15]. Second, we identified SNPs showing strong associations with environmental variables based on a latent factor mixed-effect model (LFMM) [16].

Finally we performed genome-wide association mapping (GWAS) on the timing of bud set, our target adaptive trait, using GEMMA (Fig. 2a, [17]). SNPs identified as significant (false discovery rate<0.05) by the three methods showed a large degree of overlap (Additional file3: Figure S3) and for subsequent analyses we consider SNPs that were identified as significant by at least two of the three methods to be involved in local adaptation. 99.2% of the 910 SNPs identified by all three methods and 89.1% of the additional 705 SNPs identified by two methods were located in a single region spanning c. 700 kbp on chromosome 10 (Fig. 2a,b; Additional file3: Figure S4; Additional file4: Table S3).

**Figure 2.**
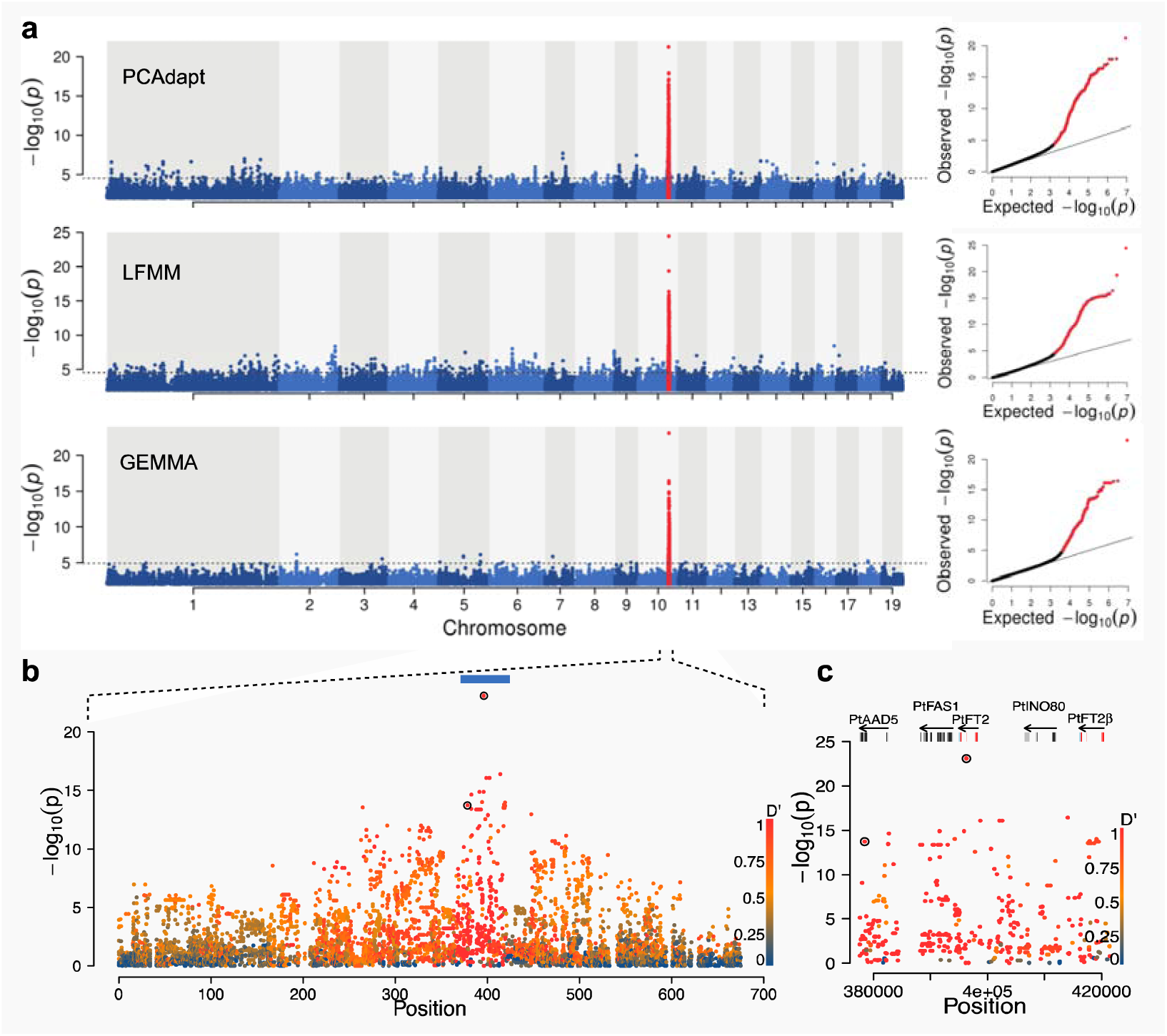
Local adaptation signals across the genome. a) Manhattan plots for SNPs associated with local adaptation using three approaches, PCAdapt, LFMM and GEMMA. The 700 kbp region surrounding *PtFT2* gene (marked in red) is identified by all methods. The dashed line represents the significance threshold for each method. Quantile-quantile plot is displayed in the right panel, with significant SNPs highlighted in red. b) Magnification of the GWAS results for the region surrounding *PtFT2* on Chr10. Individual data points are coloured according to LD with the most strongly associated SNP (Potra001246:25256). Two potential causal variants identified by CAVIAR within this region are marked by black circles. c) Close-up view of the GWAS results surrounding the two *PtFT2* homologs (red-exons, blue-UTRs) and several other genes (dark grey-exons, light grey-UTRs) on the peak region of Chr10 (depicted as blue bar in b).

SNPs associated with local adaptation displayed strong clinal patterns in allele frequencies with latitude, in stark contrast to 10,000 SNPs randomly selected from across the genome that displayed no or negligible differences among populations (Additional file3: Figure S5). The 700 kbp region on chromosome 10 encompasses 92 genes and the most strongly associated variants for all three tests are located in a region containing two *P. tremula* homologs of the *Arabidopsis FLOWERING LOCUS T*(*PtFT2*; Potra001246g10694 and an unannotated copy located c. 20 kbp upstream of *PtFT2*, tentatively named *PtFT2*β) (Fig. 2b,c). *FT* is known to be involved in controlling seasonal phenology in perennial plants [18] and has previously been implicated in regulating short-day induced growth cessation, bud set and dormancy induction in *Populus* [19-21].

We observed that *PtFT2* is conserved across *Populus* species (Additional file3: Figure S6) and although both copies of *PtFT2* appear to be expressed (Additional file3: Figure S7), the SNP showing the strongest signal of local adaptation across all three methods (Potra001246:25256) was located in the third intron of the previously annotated copy of *PtFT2* (Potra001246g10694) (Fig. 2c). This SNP explain 65% of the observed genetic variation in the timing of bud set across years and sites. Furthermore, it was identified as having highest probability of being the causal variant within the 700 kbp region by CAVIAR [22](Fig. 2b,c), a fine-mapping method that accounts for linkage disequilibrium (LD) and effect sizes to rank potential causal variants. Another potentially causal SNP (Potra001246:43095) in this region is in strong LD with Potra001246:25256 (Fig. 2c). Therefore, we identify *PtFT2* as a candidate gene, and henceforth, we refer to the entire ~700 kbp region centered on *PtFT2* as the *PtFT2* locus. We note, however, that this region potentially harbours many SNPs that could individually contribute to bud set and hence are involved in local adaptation.

### Evidence of selective sweep

In order to gain further insight into the evolutionary history of the *PtFT2* locus, we performed several haplotype-based tests to examine the presence of recent positive selection in this region. We calculated the standardized integrated haplotype score (iHS) [23] for all SNPs (8,570 SNPs where information of ancestral or derived states was available) located in the 700-kbp region (Fig. 3a). Positive selection signals, revealed by |iHS| >2.0, were observed for 20.6% of all tested SNPs. We found that the region surrounding *PtFT2* contained the highest concentration of significant hits by the iHS test across the genome (Fig. 3b), confirming that *PtFT2* locus as the strongest candidate for positive selection in the Swedish populations of *P. tremula*. Similar results were found when the number of segregating sites by length (nSL) [24], which has proven sensitive for detecting incomplete selective sweeps, was calculated for these same loci (Additional file3: Figure S8). We further performed extended haplotype homozygosity (EHH, [25]) test, centering on the most strongly associated SNP (Potra001246:25256), to explore the extent of haplotype homozygosity around the selected region. The core haplotype carrying the derived allele (G) had elevated EHH and exhibited long-range LD relative to haplotypes carrying the ancestral allele (T) (Fig. 3d). Also, haplotypes carrying the derived allele were longer than those carrying the ancestral allele (Fig. 3e). Notably, the derived allele with high EHH is largely restricted to the four high-latitude populations and almost absent in the southernmost populations (Figure 3c), implying that *PtFT2* locus has likely been subjected to geographically restricted selective sweeps.

**Figure 3.**
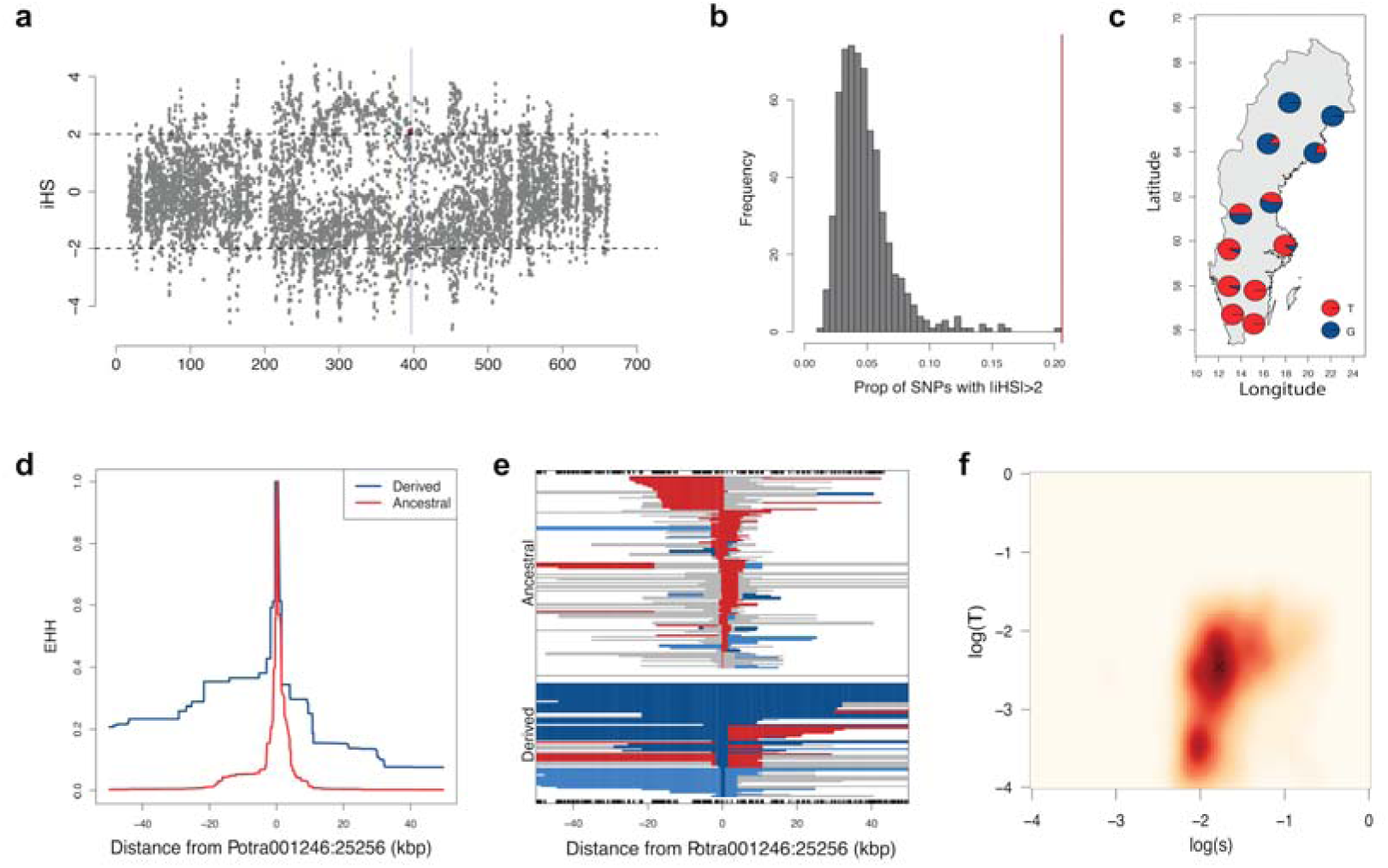
Evidence of positive selection centered on the *PtFT2* locus. a) Patterns of normalized iHS scores (y-axes) across the ~700 kbp genomic region (x-axis) around the *PtFT2* gene (vertical light grey bar). The dashed horizontal lines indicate the threshold of positive selection signal (|iHS| > 2). The red dot indicates the SNP (Potra001246:25256) showing the strongest signal of local adaptation. (b) a high concentration of significant |iHS| signals was found in the ~700 kbp region surrounding *PtFT2* (marked as red line) compared to the genome-wide distribution (based on dividing the genome into non-overlapping windows of 700 kbp). c) Allele frequencies of the most strongly associated SNP Potra001246:25256 for the twelve original populations of the SwAsp collection. d) the decay of extended haplotype homozygosity (EHH) of the derived (blue) and ancestral (red) alleles for the SNP Potra001246:25256. e) The extent of the three most common haplotypes at Potra001246:25256. Rare recombinant haplotypes were pooled and are displayed in grey. f) Joint inference of allele age and selection coefficient for the region surrounding *PtFT2.*

To further understand the evolution of functional differences between northern and southern *PtFT2* alleles, we examined the patterns of genetic variation at the *PtFT2* locus separately for South (pop 1-6), Mid (pop 7-8) and North (pop 9-12) populations. First, we found that the nucleotide diversity at the *PtFT2* locus was significantly below the genome-wide averages in all groups of populations (Fig. 4a,b; Additional file5: Table S4), which was consistent with the expectation under a selective sweep [26]. In particular, northern populations were observed to have a much stronger reduction of genetic diversity relative to other populations (Fig. 4a,b). Additionally, the level of genetic differentiation among populations was very high at *PtFT2* locus compared with genomic background, especially between southern and northern populations (Fig. 4c,d; Additional file5: Table S4). Furthermore, high H12 but low H2/H1 statistics [27] was only observed in northern populations (Fig. 4e,f,g,h; Additional file5: Table S4), providing a clear indication of a single adaptive haplotype that has risen to high frequency among these populations (Additional file3: Figure S9). Finally, we performed a composite-likelihood based (CLR) test and separately evaluated the evidence of positive selection in different groups of populations. Also, as expected for selective sweep, a distorted site frequency spectrum with an excess of rare and high frequency derived variants near the *PtFT2* locus was only found in northern populations (Fig. 4i,j; Additional file5: Table S4). Overall, all these findings provide strong evidence for the occurrence of a recent geographically restricted selective sweep in northern-most Swedish populations of *P. tremula*.

**Figure 4.**
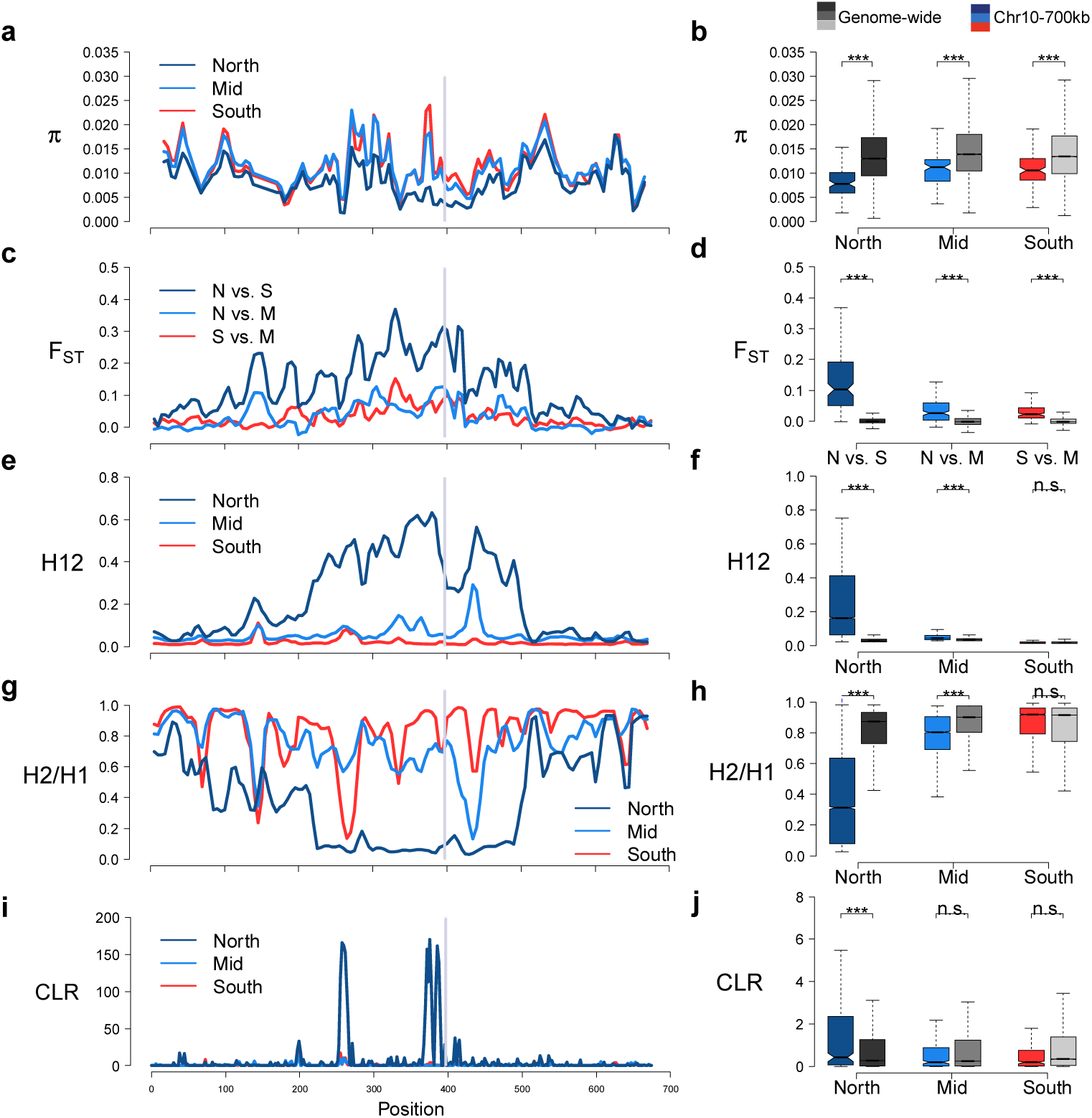
Geographically restricted selective sweep in northernmost populations. (left panels) A magnified view of different summary statistics that are sensitive to the effects of a selective sweep for the ~700 kbp region surrounding *PtFT2*. The grey bar marks the location of the *PtFT2* gene. (right panels) Comparison of these statistics between the *PtFT2* region (colored boxplot) and the genome-wide averages (grey boxplot). Statistics were calculated separately for individuals from southern (population 1-6), middle (populations 7-8) and northern (populations 9-12) in Sweden. a,b) Nucleotide diversity, π c,d) Genetic differentiation, *F*_ST_, e,f) H12, g,h) H2/H1, i,j) composite likelihood ratio (CLR) test for the presence of a selective sweep.

The observation of a single adaptive haplotype rising to high frequency in high-latitude populations (Fig. 4; Additional file3: Figure S9) is consistent with a hard selective sweep pattern, where adaptation can result either from a *de novo* mutation or from a low frequency standing variant that was already present in the population prior to the onset of selection (single origin soft sweep, cf. [28]). Assuming that the beneficial allele present in northern populations has been driven to fixation by a hard selective sweep, we used an Approximate Bayesian Computation (ABC) method [29] to jointly estimate the age and strength of selection acting on the northern allele. The results (Fig. 3f) point to a recent origin of the northern allele (T=18952 years, 95% credible interval 719 - 114122 years) and that selection during the sweep has been relatively strong (s=0.016, 95% credible interval 0.006 - 0.192). This suggests that the adaptive event that occurred in northern-most populations of *P. tremula* most likely represents an evolutionary response to the harsher environmental conditions experienced by these populations during the post-glacial colonization of northern Scandinavia.

### PtFT2 regulates the timing of bud set

Although the extensive LD in the immediate vicinity of the *PtFT*2 locus (Figure 2b) makes it hard to identify the true causal SNP(s) that are involved in mediating natural variation in bud set, we found that the significantly associated SNPs are overall enriched in non-coding regions located in and around genes and show a deficit in intergenic regions (Additional file3: Figure S10; Additional file4: Table S3). One possible way that functional variation is mediated by these SNPs is thus by altering expression patterns of related genes across the latitudinal gradient. To further assess the possibility that patterns of *PtFT*2 expression is involved in mediating local adaptation, we selected two southern genotypes and two northern genotypes for greenhouse and field experiments in order to test whether *PtFT2* expression regulates the timing of growth cessation and bud set. In greenhouse experiments, we found that the two northern genotypes showed rapid growth cessation and bud set following a shift from long (23hr day length) to short day (19hr day length) conditions whereas the two southern genotypes continued active growth under the same conditions (Fig. 5a). Analyses of *PtFT*2 gene expression in these genotypes show a strong down-regulation of *PtFT*2 in the northern genotypes in conjunction with growth cessation and bud set (Fig. 5b; Additional file6: Table S5). Similarly, under field conditions we observe that northern genotypes also show lower expression of *PtFT*2 even at a time point when all genotypes were actively growing (Fig. 5c).

**Figure 5.**
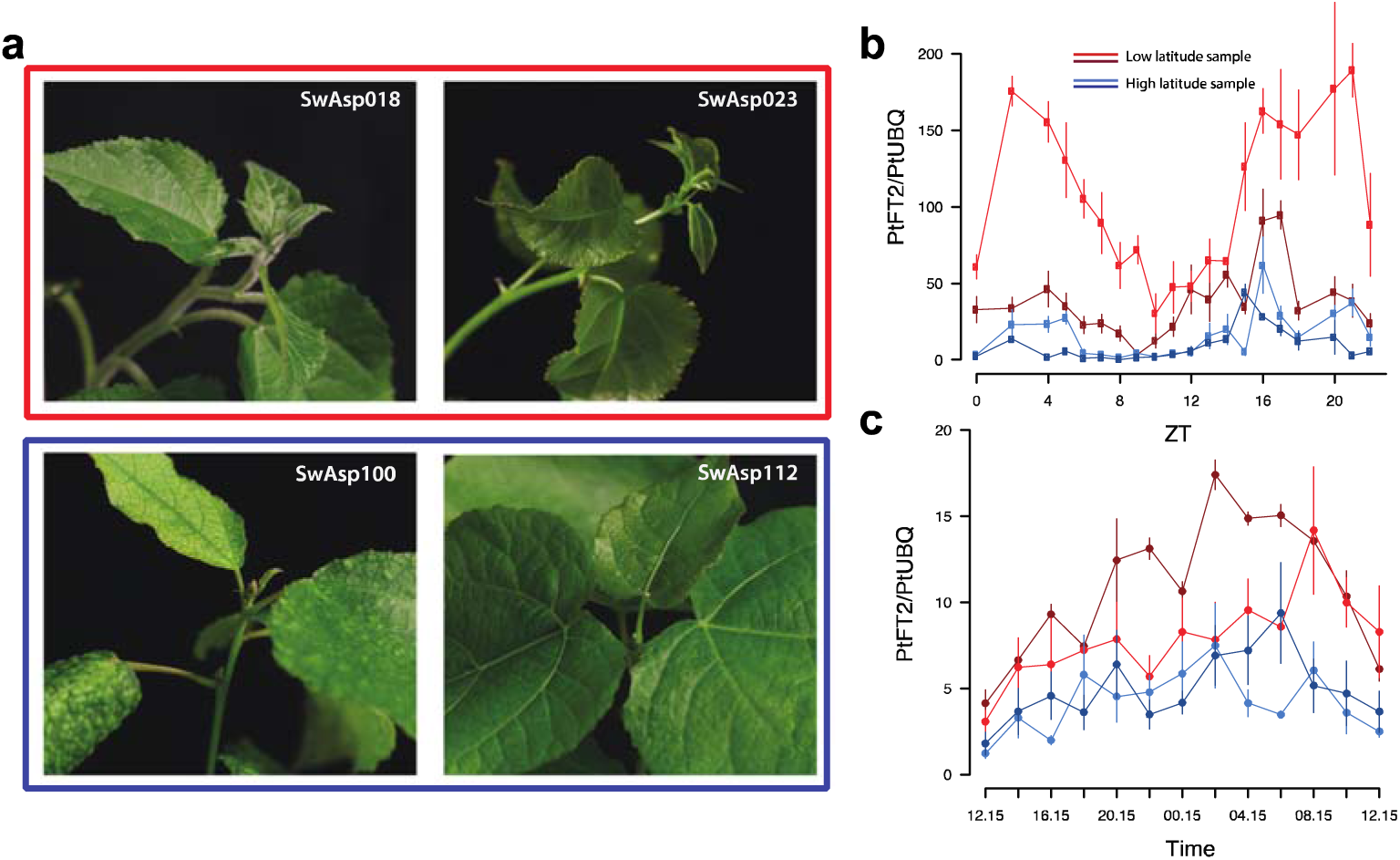
*PtFT2* expression affects short-day induced growth cessation and bud set in *P. tremula.* a) Bud set phenotype under 19hr day-length conditions. Two southern clones (SwAsp 018, Ronneby, latitude 56.2 ˚N; SwAsp 023, Vårgårda, latitudes 58 ˚N) and two northern clones (SwAsp 100, Umeå, latitude 63.9 ˚N; SwAsp 112, Luleå, latitudes 65.7 ˚N) were chosen to be analyzed. Trees were grown under 23hr day-length for one month and then shifted to 19hr day-length, Photos were taken one month after the shift to 19hr day-length. b) Dynamic expression analysis of *PtFT2* in two southern clones (red) and two northern clones (blue) from the greenhouse experiment (same clone number as in Fig 3A). Samples for RT-PCR were taken two weeks after the trees were shifted to 19hr day-length. c) Dynamic expression analysis of *PtFT2* in two southern clones (red) and two northern clones (blue) from common garden experiment. Samples were collected in the Sävar common garden in early July 2014.

Furthermore, down-regulation of the *PtFT*2 expression using RNAi to approximately 20% of wild type levels accelerates bud set by c. 23 days, a difference that is comparable to the differences we observe between the most extreme phenotypes in our field-collected trees (Fig. 6). For instance, wild-collected trees carrying the derived G allele in homozygous form for the most strongly associated SNP in *PtFT2* (Potra001246:25256) set bud on average 28 days earlier than those homozygous for the ancestral T allele, with the derived G allele showing partial dominance (Fig. 6a). The RNAi experiment thus provides additional evidence that differences in gene expression of *PtFT2* are involved in mediating the phenotypic differences we observe in bud set between northern and southern genotypes.

**Figure 6.**
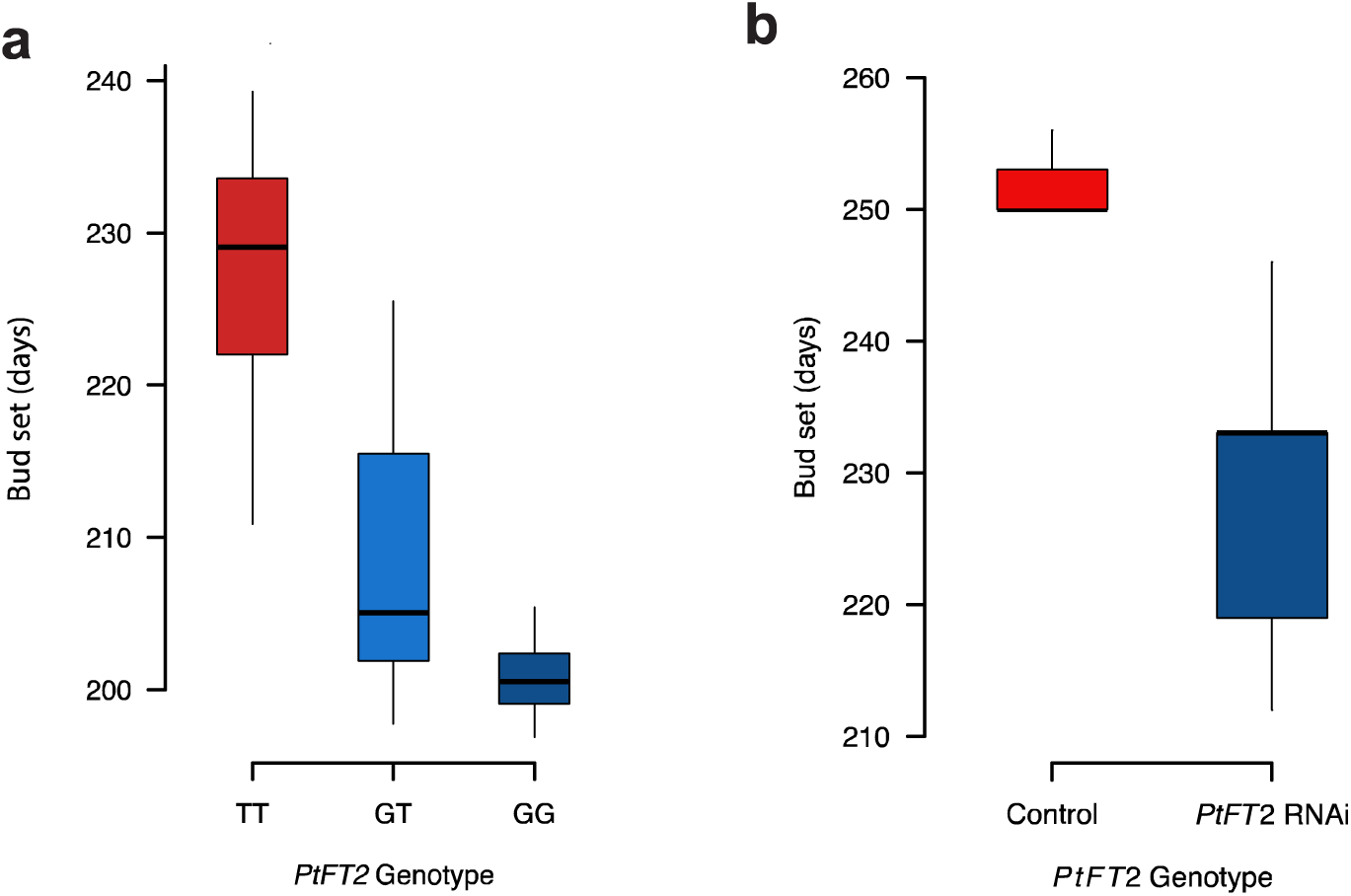
Phenotypic effects of *PtFT2*. a) Genotypic means of the timing of bud set for the three genotypes of the *PtFT2* SNP (Potra001246:25256) that displays the strongest signal of local adaptation identified by all three methods as shown in Fig. 2a. The effect displayed is the mean time to bud set of genotypes after correcting for the effects of common garden site, year and block. b) The average timing of bud set for wild type control lines and transgenic *PtFT2* lines in the field experiments at Våxtorp.

## Discussion

To date, only a small number of candidate genes have been used to identify potential loci linked to traits involved in local adaptation in *P. tremula* [11,30,31]. Here we have substantially expanded our earlier studies by utilising data from whole genome re-sequencing to local environmental variables and phenotypic variation in a key adaptive life history trait in order to investigate the genomic basis of local adaptation in *P. tremula*. Extremely weak genetic structure is found across Swedish populations along the latitudinal gradient, indicating that there is extensive gene flow likely mediated by wind dispersal of both seeds and pollen among these populations. We identify a locus, centered on *PtFT*2, that has a major effect on phenotypic variation in bud set and that has played a key role in the rapid adaptation of *P. tremula* to northern environments. This region has been subject to recent and strong selective sweep that is regionally restricted to the northernmost populations. This selective event has likely been driven by adaptation in response to the substantially shorter growing seasons that *P. tremula* has encountered at northern latitudes during the postglacial colonization of northern Scandinavia following the last glaciation.

The likely target of the selective sweep, *PtFT2*, is a *P. tremula* homolog of the *Arabidopsis FT* gene that plays a central and widely conserved role in day length perception and seasonal regulation of photoperiodic responses 2[32]. In *Populus* the *FT* gene is represented by two functionally diverged paralogs where *PtFTl* has retained the function of reproductive initiation whereas *PtFT2* acts to maintain growth and prevent bud set [19,20]. We observe that differences in *PtFT2* gene expression between genotypes from southern and northern populations are associated with the timing of bud set in response to variable day lengths in different environments (Figure5b,c). Transgenic down-regulation of *PtFT2*, under field conditions, yields a phenotype that closely mimics variation found in our wild collected trees, further implicating that non-coding regulatory variation in or around *PtFT2* likely mediate local adaptation in bud set by altering the level and timing of *PtFT2* expression.

The *FT* gene has repeatedly gone through duplications and functional diversifications in many plants, and variation within and around these *FT-like* genes are involved in mediating adaptive responses to photoperiod changes and altering overall fitness in a wide range of plant species [33]. For example, in *Arabidopsis thaliana*, naturally occurring variation in the promoter of *FT* control variation in flowering time by altering the timing and level of *FT* expression [34,35], and in rice (*Oryza sati*va) non-coding variation in the promoter of *RFT*1 results in reduced expression of *RFT*1 and a corresponding delay in flowering [36]. Similarly, in soybean (*Glycine max*) a weakly expressed allele at the *FT*2a locus delays flowering compared to the wild type allele. While the two soybean *FT*2a alleles have identical coding regions they differ at a number of sites in the promoter and intron regions, including the insertion of a *TY*1/*copia*-like retrotransposon into the first intron of the weakly expressed allele[37]. *FT-like* genes thus emerge as an evolutionary hotspot for regulating seasonal patterns in diverse annual and perennial species [38,39]. This is further illustrated by the pivotal role of *FT* in the photoperiodic pathway where *FT* serves as a nexus for integrating complex day-length sensing information and triggering cell division at the flower meristem in *Arapbidopsis* or at the apical meristem in *Populus* [18]. Given the position of *FT* within the regulatory network, evolutionary changes in *FT* are supposed to have minimal pleiotropic effects and this can help clarify why *FT*, rather than other genes in the network, has acted as a hotspot gene which has repeatedly accumulated substantial evolutionary relevant mutations [38,40] (Additional file3: Figure S11). A study in the related species *Populus trichocarpa* also identified non-coding variation of *PtFT2*, a SNP in the second intron, that were associated with naturally occurring variation in bud set [21]. Although the exact causal mutations differ, this demonstrates that parallel adaptive changes in the timing of bud set between *P. tremula* and *P. trichocarpa*, two species that diverged more than 7 million years ago and that occur on different continents, has involved changes in the same orthologous gene.

Our findings additionally provide empirical evidence supporting recent theoretical predictions that local adaptation in the face of high gene flow tends to favor few loci of large-effect rather than many loci of small effect [3]. This is because large-effect loci are more likely to establish and persist over longer time scales as they are able to resist the homogenizing effect of migration [3]. In contrast, small-effect loci are prone to swamping and only transiently contribute to local adaptation [41]. The distribution of number and effect-size for variants controlling adaptive trait is therefore expected to shift to few large-effect loci under persistent migration-selection balance [3] compared with models from isolated populations [42]. Multiple mechanisms can give rise to the characteristic pattern in *P. tremula* where a single locus explains most of the variation for a key life history trait and facilitates rapid adaptation. First, the presence of genomic rearrangements, such as chromosomal inversions, that suppress recombination can be favoured by natural selection and cause the clustering of SNPs associated with local adaptation at the *PtFT2* locus [43,44]. However, in contrast to expectations from the presence of an inversion, we did not observe blocks of elevated LD around the *PtFT2* locus (Additional file3: Figure S12). LD in this region decays rapidly and falls to background levels within a few thousand bases, similar to what is seen in other regions genome-wide (Additional file3: Figure S12a). This indicates that frequent recombination has occurred in this region and that the clustering of SNPs involved in local adaptation most likely arose from a selective sweep instead of an inversion [45]. Nonetheless, owing to the limited ability to detect inversions using short-insert paired reads, future characterization of structural variation across the genome is clearly required to determine whether genomic rearrangements are involved in mediating signals of adaptation in the *Populus* genome. Second, the establishment probability of additional adaptive mutations can be increased in the vicinity of a locus undergoing strong divergent selection, leading to a genomic architecture where multiple, tightly linked loci are controlling an adaptive trait [46]. However, recent theoretical work has shown that the conditions for such establishment of *de novo* linked beneficial mutations are rather restrictive [47]. Instead, another potentially more important mechanism for the formation of ‘genomic islands’ of strong genetic differentiation is via secondary contact and the erosion of pre-existing genetic divergence, which is a process that can be very rapid, especially compared to the alternative scenario that involves the fixation of novel mutations [47]. This mechanism provides a tantalizing hypothesis for *P. tremula* where earlier studies have established the existence of a hybrid zone between divergent post-glacial lineages in Scandinavia [14]. The selective sweep at *PtFT2* is geographically restricted and likely occurred prior to secondary contact. Therefore, the large genomic ‘island’ of divergence that we observe surrounding the *PtFT2* locus is a strong candidate for having evolved via erosion following secondary contact.

### Conclusions

Our study of phenotypic and genetic variation in *P. tremula* across a latitudinal gradient in Sweden suggests that a strong and recent selective sweep has occurred in the northernmost populations following the last glaciation. Northern and southern populations have experienced high rates of gene flow following secondary contact [14], resulting in very low levels of genome-wide genetic differentiation across the latitudinal gradient. However, the region surrounding the *PtFT2* gene differs markedly from the genome-wide background, showing strong genetic differentiation between northern and southern populations, a very pronounced haplotype structure spanning nearly 700 kbp and where segregating mutations show strong associations with naturally occurring variation of bud set. Our results suggest a scenario in which natural selection is actively maintaining alternate alleles across the latitudinal gradient in the face of high levels of gene flow. This adaptation has arisen and been driven to fixation during the post-glacial colonization of northern Scandinavia in response to the substantially shorter growing seasons that are characteristic of northern latitudes. Although the core photoperiod pathway is largely conserved in plants [32], functional diversification of *FT* has repeatedly occurred in many plant species [33]. Given the central role of *FT* as a key integrator of diverse environmental signals, it is perhaps not surprising that *FT* is acting like an evolutionary hotspot for rapid adaptation to changing environmental conditions and that these adaptations are mediated through *cis*-regulatory changes. *FT* thus appears to serve as evolutionary ‘master switch’ for adaptive life history variation, similar to what have been seen for a few other loci in plants, such *FLC* [48], *FRI*[49] and *DOG*1 [50,51].

## Materials and Methods

### Sample collection and sequencing

We collected material from all available trees in the Swedish Aspen (SwAsp), which consists of 116 individuals collected from 12 different locations spanning the distribution range in Sweden [12] (Fig. 1a). Leaf material was sampled from one clonal replicate of each individual growing at a common garden experiment located in Sävar, northern Sweden. Total genomic DNA for each individual was extracted from frozen leaf tissue using the DNeasy plant mini prep kit (QIAGEN, Valencia, USA). Paired-end sequencing libraries with an average insert size of 650 bp were constructed for all samples according to the Illumina manufacturer’s instructions. Whole genome sequencing and base calling were performed on the Illumina HiSeq 2000 platform for all individuals to a mean, per-sample depth of approximately 30× at the Science for Life Laboratory, Stockholm, Sweden.

### Sequence quality checking, read mapping and post-mapping filtering

A total of 103 SwAsp individuals were successfully sequenced. Prior to read mapping, we used Trimmomatic v0.30 [52] to identify reads with adapter contamination and to trim adapter sequences from reads. After checking the quality of the raw sequencing data using FastQC (http://www.bioinformatics.bbsrc.ac.uk/proiects/fastqc/), the quality of sequencing reads was found to drop towards the ends of reads (Additional file3: Figure S13). We therefore used Trimmomatic v0.30 to trim bases from both ends of the reads if their qualities were lower than 20. Reads shorter than 36 bases after trimming were discarded completely.

After quality control, all high-quality reads were mapped to a *de novo* assembly of the *P. tremula* genome (available at http://popgenie.org; [53]) using the BWA-MEM algorithm with default parameters using bwa-0.7.10 [54]. We used MarkDuplicates methods from the Picard packages (http://picard.sourceforge.net) to correct for the artifacts of PCR duplication by only keeping one read or read-pair with the highest summed base quality among those of identical external coordinates and/or same insert lengths. Alignments of all paired-end and single-end reads for each sample were then merged using SAMtools 0.1.19 [55]. Sequencing reads in the vicinity of insertions and deletions (indels) were globally realigned using the RealignerTargetCreator and IndelRealigner in the Genome Analysis Toolkit (GATK v3.2.2) [56]. To minimize the influence of mapping bias, we further discarded the following site types: (i) sites with extremely low (<400× across all samples, i.e. less than an average of 4× per sample) or extremely high coverage (>4500×, or approximately twice the mean depth at variant sites) across all samples after investigating the coverage distribution empirically; (ii) sites with a high number of reads (>200×, that is on average more than two reads per sample) with mapping score equaling zero; (iii) sites located within repetitive sequences as identified using RepeatMasker [57]; (iv) sites that were in genomic scaffolds with a length shorter than 2 kbp.

### SNP and genotype calling

SNP calling in each sample was performed using the GATK HaplotypeCaller, and GenotypeGVCFs were then used to perform the multi-sample joint aggregation, regenotyping and re-annotation of the newly merged records among all samples. We performed several filtering steps to minimize SNP calling bias and to retain only high-quality SNPs: (i) Remove SNPs at sites not passing all previous filtering criteria; (ii) Retain only bi-allelic SNPs with a distance of more than 5 bp away from any indels; (iii) Remove SNPs for which the available information derived from <70% of the sampled individuals after treating genotypes with quality score (GQ) lower than 10 as missing; (iv) Remove SNPs with an excess of heterozygotes and deviates from Hardy-Weinberg Equilibrium test (*P*-value <1e-8). After all steps of filtering, a total of 4,425,109 SNPs with minor allele frequency higher than 5% were left for downstream analysis. Finally, the effect of each SNP was annotated using SnpEff version 3.6 [58] based on gene models from the *P. tremula* reference genome (available at http://popgenie.org; [53]), and the most deleterious effect was selected if multiple effects occurred for the same SNP using a custom Perl script.

### Relatedness, population structure and isolation-by-distance

To identify closely related individuals and to infer population structure among the sampled individuals, we discarded SNPs with missing rate >10%, MAF < 5% and that failed the Hardy-Weinberg equilibrium test (*P* <1×10^−6^) after all filtering steps as shown above. We also generated LD-trimmed SNP sets by removing one SNP from each pair of SNPs when the correlation coefficients (*r*^2^) between SNPs exceed 0.2 in blocks of 50 SNPs using PLINK v1.9 [59]. This yielded 217,489 independent SNPs that were retained for downstream analyses of population structure. First, we used PLINK v1.9 to estimate identity-by-state (IBS) scores among pairs of all individuals. Nine individuals were excluded from further analyses due to their high pairwise genetic similarity with another sampled individual (IBS>0.8), leaving a total of 94 ‘unrelated’ individuals for all subsequent analyses (Additional file3: Figure S14). Then, we used the smartpca program in EIGENSOFT v5.0 [13] to perform the principal component analysis (PCA) on the reduced set of genome-wide independent SNPs. A Tracey-Widom test, implemented in the program twstats in EIGENSOFT v5.0, was used to determine the significance level of the eigenvectors. Finally, isolation by distance (IBD) analysis was computed based on the pairwise comparison of the genetic and geographic distances between populations. We calculated the population differentiation coefficient (*F*_ST_) [60] for each pair of the twelve populations using VCFtools v0.1.12b [61]. The relationship between genetic distance measured as *F*_ST_/(1-*F*_ST_) and geographic distance (km) was evaluated using Mantel tests in the R package “vegan” [62], and the significance of the correlation was estimated based on 9999 permutations.

### Screening for SNPs associated with local adaptation

We used three conceptually different approaches to test for genome-wide signatures of local adaptation. First, we detected candidate SNPs involved in local adaptation using the principal component analysis as implemented in PCAdapt [63]. PCAdapt examines the correlations (measured as the squared loadings *ρ*^2^_jk_, which is the squared correlation between the *j*th SNP and the *k*th principal component) between genetic variants and specific principal components (PCs) without any prior definition of populations. As only the first principal component was significant from the PC analysis (see Results), we only estimated the squared loadings *ρ*^2^_j1_ with PC1 to identify SNPs involved in local adaptation. Our results showed that most outlier SNPs that were highly correlated with the first population structure PC also had high *F*_ST_ values between populations (Additional file3: Figure S15). Assuming a chi-square distribution for the squared loadings *ρ*^2^_j1_, as suggested by [63], we used PCAdapt to compute *P*-values for all SNPs, and then calculated the false discovery rate (FDR) using the method of [64] to generate a list of candidate SNPs showing significant associations to population structure. Only SNPs with FDR < 5% were retained as those significantly involved in local adaptation.

Second, we tested for the presence of candidate SNPs that exhibited high correlations with environmental gradients. To do this, a total of 39 environmental variables were analysed (Additional file7: Table S6). Precipitation and temperature values were retrieved from WorldClim version 1 [65]. Sunshine hours, photosynthetically active radiation and UV radiation were obtained using the STRÅNG data model at the Swedish Meteorological and Hydrological Institute (SMHI) (http://strang.smhi.se). Values were collected from the years 2002-2012 for the original sample coordinates of each SwAsp individual and the average values over years were then calculated. The environmental variables include latitude, longitude, altitude, the number of days with temperatures higher than 5 °C, UV irradiance, the photosynthetic photon flux density (PPFD), sunshine duration, monthly and annual average precipitation and temperature. Due to the high degree of correlation among these environmental variables (Additional file3: Figure S16a), we performed a PCA on these variables using the ‘prcomp’ function in R to identify PCs that best summarized the range of environmental variation. The first environmental PC, which explained > 60% of the total variance (Additional file3: Figure S16b,c) and had the strongest loadings for the length of growing season (Additional file3: Figure S16d), was kept to represent our target environmental variable for further analyses. We then used a latent factor mixed-effect model (LFMM) implemented in the package LEA in R [66] to investigate associations between SNPs and the first environmental PC while simultaneously accounting for population structure by introducing unobserved latent factors into the model [16]. Due to the weak population structure found in the SwAsp collection (see Results), we ran the LEA function *lfmm* with the number of latent factors (*K*) ranging from one to three, using 5000 iterations as burn-in followed by 10,000 iterations to compute LFMM parameters for all SNPs. This was performed five times for each value of *K*, and we observed identical results both across different values of *K* and across independent runs within each value of *K* (data not shown). We only showed the results using *K*=2 to account for the background population structure.LFMM outliers were detected as those SNPs with FDR < 0.05 after using the method of [64] to account for multiple testing.

Third, we obtained previously published measurements of the timing of bud set, which is a highly heritable trait that shows strong adaptive differentiation along the latitudinal gradient [10,31]. To measure phenotypic traits, all SwAsp individuals have previously been clonally replicated (four ramets per individual) and planted at two common garden sites in 2004 (Sävar, 63°N, and Ekebo, 56°N) (Fig. 1a). The common garden setup is described in detail in [12]. The timing of bud set was scored twice weekly starting from mid-July and continuing until all trees had set terminal buds. Bud set measurements were scored in three consecutive years, from 2005 to 2007, in both common gardens [10]. A severe drought in Sävar caused most of the trees to set bud prematurely in 2006, and we therefore excluded data from Sävar in 2006 in all downstream analyses (see [31] for further discussion). We combined data on bud set from the two common garden sites and years by predicting genetic values with best linear unbiased prediction (BLUP) for all individuals. The ASReml [67] was used to fit Equation 1 to the data for calculating BLUP using restricted maximum-likelihood techniques.

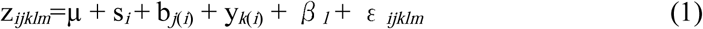

where *Z_ijklm_* is the phenotype of the *m*th individual in the *j*th block in the *k*th year of the *l*th clone from the *i*th site. In Equation 1, μ denotes the grand mean and *ε _ijklm_*is the residual term. The clone (*β _l_*, BLUP) and residual term ( *ε _ijklm_*) were modeled as random effects, whereas the site (s_*i*_,), site/block (b_*j*(*i*)_) and site/year (y_*k*(*i*)_) were treated as fixed effects. The genetic value of each individual was then used as the dependent trait in an univariate linear mixed model for SNP-trait association analyses performed with GEMMA [17]. This method takes relatedness among samples into account through the use of a kinship matrix. The mixed model approach implemented in GEMMA has been shown to outperform methods that try to correct for population structure by including it as a fixed effect in the GWAS analyses [68]. Given the extremely weak population structure we observe in our GWAS population (see Results) we did not pursue any further corrections for population structure in the association analyses as this likely would severely reduce our power to detect significant associations. As described previously, we used a FDR < 5% [64] to control for the multiple testing across the 4,425,109 SNPs.

### Genotype imputation

For some haplotype-based selection tests, imputed and phased data sets were needed. We therefore used BEAGLE v4.1 [69] to perform imputation and haplotype phasing on genotypes of 94 individuals with default parameters. Before performing genotype imputation, we first used Chromosemble from the Satsuma packages [70] to order and orient the scaffolds of the *P. tremula* assembly to 19 pseudo-chromosomes according to synteny with the *P. trichocarpa* genome. We then performed pairwise genome alignment between scaffolds of *P. tremula* and the 19 pseudochromosomes using the BLAST algorithm (*E*-value cutoff of 1e-50), and finally, more than 99% of the SNPs (4,397,537 out of 4,425,109) were anchored on the 19 pseudochromosomes.

To test for the accuracy of imputation, and its relationship with the MAF cutoff and the missing rate of genotypes in our dataset, we selected 346,821 SNPs with a rate of missing genotypes lower than 10% from the pseudo-chromosome 2 (~32.6 Mb) for the simulation analysis. We randomly masked out varying proportions (5-50%) of SNPs, which were treated as missing. BEAGLE v 4.1 was then used to impute genotypes at the masked positions. We found high imputation accuracy (>0.97) across a wide range of MAF when rates of missing genotypes were less than 30% (Additional file3: Figure S17), suggesting imputation and phasing by BEAGLE should not bias the accuracy of our results. We therefore phased and imputed genotypes of the SNPs anchored on pseudo-chromosomes using BEAGLE v 4.1.

### Estimation of ancestral states for all SNPs

Since the ancestral states of SNPs are usually used for selection detection, for each SNP, we classified alleles as either ancestral or derived on the basis of comparisons with two outgroup species: *P. tremuloides* and *P. trichocarpa*. We obtained publicly available short read Illumina data for one *P. tremuloides* (SRA ID: SRR2749867) and one *P. trichocarpa* (SRA ID: SRR1571343) individual from the NCBI Sequence Read Archive (SRA) [71]. We individually aligned the reads from these two samples to the *de novo P. tremula* assembly (Potra v1.1, available at PopGenIE.org) and used UnifiedGenotyper in GATK to call SNPs at all sites (-- output_mode EMIT_ALL_SITES). For each SNP, two procedures were performed to define their ancestral states: (1) because *P. trichocarpa* is more distantly related to *P. tremula* compared to *P. tremuloides* [72] and from our previous study there were less than 1% polymorphic sites shared between *P. tremula* and *P. trichocarpa* [71], we inferred the ancestral state as the *P. trichocarpa* allele at sites where the *P. trichocarpa* individual was homozygous and matched one of the *P. tremula* alleles; Otherwise, (2) we inferred the ancestral state as the *P. tremuloides* allele at sites where the *P. tremuloides* individual was homozygous and matched one of the *P. tremula* alleles. If the above two requirements were not met, the ancestral state was defined as missing. In total, we obtained information of ancestral states for 96.3% of all SNPs.

### Anchoring and orientation of SNPs associated with local adaptation to a single region on chromosome 10

As we found that a large majority of significant SNPs (>90%) detected by at least two of the three methods (PCAdapt, LFMM, and GEMMA) were clustered in a single genomic region on pseudo-chromosome 10, we performed several further steps to refine the anchoring and orientation of these SNPs within this region. First, we used ColaAlignSatsuma from the Satsuma packages [70] to align the genomes of *P. tremula* and *P. trichocarpa* using default settings. The output was then converted and filtered into GBrowse synteny compatible format that was available at http://popgenie.org [53]. Based on the alignment of the two genomes, 15 scaffolds from the *P. tremula* assembly that contain SNPs inferred to be associated with local adaptation were completely or partially mapped to a single region on chromosome 10 of *P. trichocarpa* genome (Additional file4: Table S3). We then retained only seven scaffolds that were completely mapped to the region and with length longer than 10 kbp. The seven scaffolds contained more than 95% (1465 out of 1528) of the total number of significant SNPs in the single region of chromosome 10. Lastly, according to the alignment results between the genome of *P. tremula* and *P. trichocarpa*, we reordered and re-oriented the seven scaffolds to a ~700 kbp region for all downstream selection tests (Additional file3: Figure S4).

### Linkage disequilibrium

To explore and compare patterns of LD between the ~700 kbp region on chromosome 10 and genome-wide levels, we first calculated correlations (D’ and *r*^2^) between all pairwise common SNPs (MAF>5%, 9149 SNPs) in the ~700 kbp region using PLINK 1.9 [59]. Then we used PLINK 1.9 to randomly thin the number of common SNPs across the genome to 200,000, and calculated the squared correlation coefficients (*r*^2^) between all pairs of SNPs that were within a distance of 100 kbp. The decay of LD against physical distance was estimated using nonlinear regression of pairwise *r*^2^ vs. the physical distance between sites in base pairs [73].

### Fine-mapping the causal variants using CAVIAR

We utilized CAVIAR (CAusal Variants Identification in Associated Regions, v1.0) [22] to identify the potential causal variants in the ~700 kbp region on chromosome 10. CAVIAR is a fine-mapping method that quantifies the probability of each variant in a locus to be causal and outputs a set of variants that with a predefined probability (e.g., 95% or 99%) contain all of causal variants at the locus. We created the LD structure by computing *r*^2^ between all pairwise significantly associated SNPs in the ~700 kbp region using PLINK 1.9. Marginal statistics for each significantly associated variant is the association statistics obtained from GWAS analysis by GEMMA. In our analysis, we set the causal confidence as 99% (-r 0.99) to obtain a set of causal variants that capture all the causal variants with the probability higher than 99%.

### Positive selection detection

We measured two haplotype-based tests, integrated haplotype score (iHS) [23] and the number of segregating sites by length (nS_L_) [24], to test for possible positive selection. These statistics were calculated for all SNPs with MAF higher than 0.05 and with information on ancestral state across the genome using the software `selscan` v1.1.0a [74] with its assumed default parameters. The iHS and the nS_L_ values were then normalized in frequency bins across the whole genome (we used 100 bins). To test for whether there is significant concentration of selection signals on the region surrounding the *PtFT2*, we divided the 19 pseudo-chromosomes (without the seven scaffolds around the *PtFT2* locus) into non-overlapping windows of 700 kbp and calculated the proportion of SNPs with |iHS| > 2 or with |ßnS_L_|>2 in each window. Statistical significance was assessed using the ranking of genome-wide windows, with windows having fewer than 100 SNPs being excluded.

### Population-specific selective sweeps

Several standard methods were further applied to search for signs of selective sweeps in different groups of populations: (i) pairwise nucleotide diversity (π) [75], which is expected to have a local reduction following a selective sweep, was calculated using a sliding window approach with window size of 10 kbp and moving step of 5 kbp using the software package - Analysis of Next-Generation Sequencing Data (ANGSD v0.602)[76] separately for South (pop 1-6), Mid (pop 7-8) and North (pop 9-12) populations. Only the reads with mapping quality > 30 and the bases with quality score >20 were used in the analysis. Windows with <10% of covered sites remaining from the previous filtering steps (section 2.1) were excluded; (ii) Weir and Cockerham’s *F*_ST_, which measures genetic divergence between pairs of three groups of populations, South, Mid and North, was calculated using a sliding-window approach with window size of 10 kbp and moving step of 5 kbp by VCFtools; (iii) a combination of H12 and H2/H1 [27], which measures haplotype homozygosity and can distinguish hard from soft selective sweeps, were calculated in windows of 200 SNPs (~15 kbp) for common SNPs with MAF higher than 5% separately for South, Mid and North populations. As the mean LD (*r*^2^) in *P. tremula* decays to less than 0.1 within 10 kbp (Additional file3: Figure S12a and [71]), the use of ~15 kbp windows should be large enough to differentiate the footprint of selective sweeps from those caused by neutral processes. The H12 and H2/H1 values were then averaged using a sliding window method with window size of 10 kbp and moving step of 5 kbp; (iv) a composite likelihood ratio statistic (CLR) [77], which contrasts the likelihood of the null hypothesis based on the genome-wide site frequency spectrum with the likelihood of a model where the site frequency has been altered by a recent selective sweep, was computed using `SweepFinder2` [78] separately for South, Mid and North populations. `SweepFinder2` is most efficient when information on the ancestral and derived states is available for SNPs and we therefore polarized SNPs as described above. The small fraction of SNPs (~3.7%) that could not be polarized were excluded from further analysis using `SweepFinder2`. CLRs were calculated using non-overlapping windows with a spacing of 2 kbp, and the empirical site frequency spectrum across the whole *P. tremula* genome was estimated using the −`f` option in `SweepFinder2` after including all polymorphic sites in the genome (a total of 8,007,303 SNPs). As recommended by [79] we only used sites that were polymorphic or that represented fixed substitutions in each group of populations to scan for sweeps. To determine whether there are significant differences of the above statistics between the 700 kbp region around *PtFT2* gene on chromosome 10 and genome-wide estimates, we use the same strategy to divide the genome into the windows with the same size for each test and calculated the above statistics across the genome (Results are shown in Fig. 4b,d,f,h,j and Additional file5: Table S4). Significance for the above statistical measurements was evaluated using Mann-Whitney tests.

To assess the scale of a genomic region that is affected by a selective sweep, we ran coalescent simulations modeling a selective sweep in the Northern populations. Simulations were run assuming that the selected site was located at the centre of the simulated region. Parameters for the simulations were taken from ABC calculations dating the selective sweep inferred in the North populations (as shown below).Briefly, we used a scaled population mutation rate (4N_e_μ) of 0.0081/bp, which corresponds to the average observed diversity in the North populations. Similarly we set the scaled population recombination rate (4N_e_r) to 0.0019 to match the genome-wide ratio of r/μ=0.229 in *P. tremula* [71]. Analyses of the simulated data using `SweepFinder2` showed that a single selective sweep often yields multiple significant peaks across a region spanning up to, and even exceeding, 100 kbp (95% quartile: 148221 bp; Additional file3: Figure S18).

### Dating the selective sweep in the North populations

To date the inferred selective sweep in the North populations we used the Approximate Bayesian Computation (ABC) method described in [29] to jointly estimate *s* (the strength of selection on the beneficial mutation causing the sweep) and *T* (the time since the beneficial allele fixed) assuming a model of selection from a *de novo* mutation (hard selective sweep). We simulated 5×10^5^ independent selective sweep events using the coalescent simulation program `msms` [80]. For the coalescent simulations, the ancestries of samples were traced backwards in time using standard coalescent methods and allowing for recombination. Selection was modelled at a single site by applying forward simulations, assuming additive selection so that the fitness of heterozygous and homozygous genotypes carrying the selected (derived) allele were 1 + s/2 and 1 + s, respectively. We simulated a chromosome region consisting of L=25000 sites and assumed a diploid effective population size of N_e_=92000, a mutation rate of μ=3.75×10^−8^ per base pair per generation[81], and a recombination rate of r=0.729×10^−8^ per base pair per generation. Together these parameters yielded a scaled population mutation rate equal to Θ=4N_e_μL=86.27 and a scaled population recombination rate ρ=4N_e_rL=19.76. For each simulation, values for both *s* and *T* were drawn from uniform prior distributions, log_10_(T)~U(-4,-0.5) and log10(s)~U(-4,-0.5).

### Gene expression ofPtFT2 under active growth and during growth cessation

Samples used for the expression analysis of *PtFT2* were collected from both climate chamber and the field (Sävar, 63.4°N, Umeå) conditions. For treatment in the climate chamber, two southern clones (SwAsp018, 56.2°N, Ronneby; SwAsp023, 56.2°N, Ronneby) and two northern clones (SwAsp100, 63.9°N, Umeå; SwAsp112, 65.6°N, Luleå) were selected. Plants were grown under 23h day lengths for one month and then transferred to 19h day length condition for 2 weeks before the start of sampling. Field samples were collected in the Sävar common garden in early July, 2014 and samples were taken from two southern clones (SwAsp005, 56.7°N,Simlång; SwAsp023, 56.2°N, Ronneby) and two northern clones (SwAsp100, 63.9°N, Umeå; SwAsp116, 65.6°N, Luleå). Leaves were harvested from three different clonal replicates to serve as biological repeats, flash-frozen in in liquid nitrogen and stored at -80°C until sample preparation. Samples were collected at 2h intervals for a total period of 24 h. RNA extraction for all samples was performed using a CTAB-LiCl method [82]. CDNA synthesis was performed using the iScript cDNA Synthesis Kit (BIO-RAD) according to the manufacturer’s instructions. Quantitative real-time PCR analyses were performed using a Roche LightCycler 480 II instrument, and the measurements were obtained using the relative quantification method [83]. We used primers qFT2F (5’-AGCCCAAGGCCTACAGCAGGAA-3’) and qFT2R (5’-GGGAATCTTTCTCTCATGAT-3’) for amplifying the transcript of *FT2* and qUBQF (5’-GTTGATTTTTGCTGGGAAGC-3’) and qUBQR (5’-GATCTTGGCCTTCACGTTGT-3’) for *UBQ* as the internal control. We assessed the presence of transcription of both *PtFT*2 (Potra001246g10694) and *PtFT*2βdigesting RT-PCR products with *Sac*I that distinguish the two transcripts (Additional file3: Figure S7).

### Field experiment with transgenic PtFT2 lines

Construction of the *PtFT* RNAi lines are described in detail in [19]. Transformed plants were planted together with wild type (WT) controls in a common garden at Våxtorp, Halland (latitude 56.4N, longitude 13.1E) in 2014. 18 replicates of each line were planted in a complete randomized block design together with six WT controls per block. Starting in 2015, data were collected on growth cessation, bud formation and bud set for all trees in the common garden. From early August plants were visually inspected roughly every five days and top shoots were scored according to a pre-determined scoring sheet (Additional file3: Figure S19) and classified as active growth (score 3), growth cessation (score 2), bud formation (score 1) and bud set (score 0). Scoring was continued until all plants had completely senesced in late October. Bud scoring data was converted to Julian date of bud set and analysed using the following linear model:

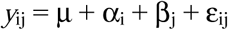

where μ is an overall mean, α_i_ is the effect of treatment *i* (where *i* is either *PtFT2* RNAi or WT) and β_j_ is the effect of block *j* and ε_ij_ are individual residual errors.

## Data availability

### Sequence and SNP data

The raw sequencing reads have been deposited in NCBI’s short-read archive, SRA, under accession number PRJNA297202. In total we identified 8,007,303 SNPs and 4,425,109 SNPs with MAF high than 5% and VCF files with SNPs (original and phased) and SNP annotations are available to download from ftp://plantgenie.org/Data/PopGenIE/Populustremula/v1.1/VCF/.

### Other data

Bud set genetic values (BLUPs) for all clones used in the GWAS is available from zendo.org (https://doi.org/10.5281/zenodo.844372).

### BASH, Perl, Python and R scripts

All scripts used for the analysis described are available in an online repository at https://github.com/parkingvarsson/PhotoperiodLocalAdaptation

## Acknowledgements

We thank Carin Olofsson for extracting DNA for all samples used in this study. STRÅNG data are obtained from the Swedish Meteorological and Hydrological Institute (SMHI), which were produced with support from the Swedish Radiation Protection Authority and the Swedish Environmental Agency. The research was funded through grants from Vetenskapsrådet, Knut and Alice Wallenbergs stiftelse and a Young Researcher Award from Umeå University to PKI. JW was supported by a scholarship from the Chinese Scholarship Council. BT is supported by the UPSC “Industrial graduate school in forest genetics, biotechnology and breeding”. NRS is supported by the Trees and Crops for the Future (TC4F) project. The authors also would like to acknowledge support from Science for Life Laboratory and the National Genomics Infrastructure (NGI) for providing assistance with massive parallel sequencing. All analyses were performed on resources provided by the Swedish National Infrastructure for Computing (SNIC) at Uppsala Multidisciplinary Center for Advanced Computational Science (UPPMAX) under the projects b2010014 and b2011141.

## Author contributions

JW, ON, SJ, NS and PKI conceived of and designed the experiments. JW, BT, AJ, BN, DGS, NS, PKI carried out all population genetic analyses. JD performed greenhouse and RT-PCR experiments. KMR and IHM collected common garden data.JW and PKI wrote the paper. All authors commented on the manuscript.

## Additional files

### Additional file1

**Table S1.** Geographical details of the 94 *Populus tremula* samples used in this study and the summary statistics of Illumina re-sequencing data per sample

### Additional file2

**Table S2.** Tracy-Widom statistics for the first three eigenvalues in PCA analysis

### Additional file3

**Figure S1-S19. Figure S1.** Genetic pairwise differentiation plotted against geographical distances between populations. **Figure S2.** Histogram of Weir & Cockerham’s *F_ST_* values among the twelve populations. **Figure S3.** Venn diagram showing the overlap of significant SNPs detected using three approaches of PCAdapt, LFMM and GEMMA. **Figure S4.** Magnified view of negative log_10_-transformed *P* values calculated from three approaches, PCAdapt, LFMM and GEMMA (from top to bottom) for all SNPs in the seven adjacent scaffolds (with length > 10 kbp, shown as boxes with alternating shades), which were anchored to a single region (~700 kbp) on chromosome 10 based on the genome alignment between *P. tremula* and *P. trichocarpa*. The significant SNPs (at false discovery rate of *Q*-value <0.05) identified by all three approaches are denoted by red dots, and those identified by two of the three approaches are denoted by pink dots. The light blue dots represent those nonsignificant SNPs. Dotted horizontal line represents the genome-wide average value of -log_10_(*P*) calculated by each method. The genomic locations of *PtFT2* gene within this region are shaded as a grey bar. The seven scaffolds from left to right are Potra000799, Potra000908, Potra000342, Potra001246, Potra004002, Potra003230, Potra000530, respectively. **Figure S5.** Derived- and ancestral- allele frequency plotted against population for significant SNPs involved in local adaptation on chromosome 10 (a) and for 10,000 randomly selected SNPs from the genome (b). Alleles are polarized according to the signs of the Spearman’s rank correlations with the first environmental principal components (PC1), where only the derived (grey boxes) or ancestral (white boxes) allele with a positive correlation with the environmental PC1 is shown for each SNP. **Figure S6.** Plot of a pairwise alignment for genome region containing Potra001246g10694 (*PtFT2*) and Potra001246g10695 with *PtFT2*β, and the corresponding genomic region from *P. trichocarpa*, *P. deltoides* and *Salix purpurea*. Curves were calculated using default VISTA thresholds based on percentage identity (y-axis) and base pair position (x-axis), and only the regions longer than 100 bp with average conservation score above threshold of 50% were colored (exons in blue, introns and promoter in pink, and UTR in light blue). **Figure S7.** a) Nucleotide alignment between the two copies of *PtFT*2 located on scaffold Potra001246 - Potra001246g10694 and PtFT2β. The red box entitles *Sac*I a CAPS marker that distinguishes the two loci using the presence (Potra001246g10694) or absence (PtFT2β) of a SacI cut site. b) Protein alignment of - Potra001246g19694 and PtFT2β. c) Transcripts of *PtFT*2 and *PtFT*2β are distinguished by the presence /absence of a *Sac*I cut site. Top figure show restriction digests with *Sac*I of PCR products targeting *PtFT*2/*PtFT*2β using genomic DNA as a template. The bottom figures show corresponding PCRs using cDNA taken from plants growing in 18h light and 6 hr dark as a template. **Figure S8.** A high concentration of significant nSL signals was found in the ~700 kbp region around *PtFT2* gene. (a) Patterns of normalized nSL scores (y-axes) across the ~700 kbp genomic region (x-axis) around the *PtFT2* gene (shaded as grey bar). The dashed horizontal lines indicate the threshold of positive selection signal (|nSL|>2.0). The red dot indicates the SNP (Potra001246:25256) showing the strongest signal of local adaptation. We then divided the genome into 626 non-overlapping regions with size of 700 kbp (without the candidate region and regions with less than 1000 SNPs left were removed) and calculated the proportion of significant |nSL| SNPs (MAF >5%) that lie in each 700 kbp region. (b) The ~700 kbp region around *PtFT2* gene (the red lines) contained significant (empirical *P*-value <0.05) higher proportion of SNPs with signals of positive selection relative to genome-wide distribution (dark grey bars) (ranked 23^th^ among 627 regions). **Figure S9.** H12 scan for selective sweeps. (a) H12 scan in three groups of populations, South (pop 1-6, bottom), Mid (pop 7-8, middle) and North (pop 9-12, top), across the ~700 kbp region around *PtFT2* gene on chromosome 10. Each data point represents the H12 values calculated at each common SNP (minor allele frequency higher than 5%). The genomic location of *PtFT2* gene within this region is shaded as a grey bar. We picked the SNP (Potra001246:25256, red square) showing the strongest signal of local adaptation and another three randomly selected SNPs (green square) to show the haplotype frequency spectra (b-e) in each group of populations at this region (b-e, corresponding to the locations of the four SNPs from left to right). In each haplotype frequency spectra plot, the height of the first bar (light blue) in each frequency spectrum indicates the frequency of the most prevalent haplotype in each group of samples, and heights of subsequent colored bars indicate the frequency of the second, third and so on most frequent haplotypes in the samples. Grey bars indicate singletons. The values of H12 and H2/H1 are shown at the bottom of each bar plot. In northern populations, there is mainly a single haplotype dominating the haplotype spectra, indicative of hard sweeps with high H12 values but low H2/H1 values. No obvious selective sweep signals were found in either middle or southern populations. **Figure S10.** Enrichment of various functional categories in significant SNPs associated with local adaptation (red line). Grey dots show the distribution of results with 10000 bootstrap replicates. The dashed line shows the expected enrichment under the null hypothesis of no enrichment. Enrichment that is significant relative to the bootstrap method are denoted by asterisks (*P*<0.001). **Figure S11.** Signatures of local adaptation for a set of 20 candidate genes controlling phenology in *Populus*. (a-c) Distribution of negative log10-transformed *P* values calculated from three approaches, PCAdapt (a), LFMM (b) and GEMMA (c) for common SNPs (minor allele frequency>5%) within the 20 candidate genes (red lines) and all other genes across the genome (black lines). For the 20 candidate genes, the bottom and the top of the error bars represent the lowest and highest negative log_10_-transformed *P* values of each method. For the rest of genes across the genome, the bottom and the top of the error bars represent 0.5^th^ and 99.5^th^ percentiles negative log_10_-transformed *P* values. From the results, we found that except for *PtFT2* gene, it is hard to distinguish the signatures of local adaptation of all other candidate genes from the rest of genes across the genome. (d) The list of gene names (corresponding to the *Populus trichocarpa* v3 assembly and *P. tremula* v1.1 assembly) of 20 candidate genes homologous to the *Arabidopsis thaliana* phenology genes shown in a-c. **Figure S12.** (a) Decay of linkage disequilibrium (LD). The comparison of mean LD decay (estimated as *r*^2^) with physical distance between the ~700 kbp region on chromosome 10 and genome-wide average level. (b) Pairwise linkage disequilibrium (quantified using D’) among the 9149 common SNPs (minor allele frequency higher than 5%) within the ~700 kbp region on chromosome 10. **Figure S13.** Comparison of per-base sequence quality between raw and filtered sequence data in one SwAsp sample (SwAsp009) as an example. Per-base sequence quality comparison between raw paried-end sequence data (forward reads: top left and reverse reads: top right), and filtered sequence data with both forward (bottom left) and reverse (bottom middle) reads left or only single-end (bottom right) reads left. The x-axis of the BoxWhisker plot shows the position in read, and y-axis shows the quality scores. The higher the score the better the base call. The background of the plot divides the y axis into very good quality calls (green), calls of reasonable quality (orange), and calls of poor quality (red). The central red line is the median quality value, and yellow box represents the inter-quantile of quality, the upper and lower whiskers represent the 10% and 90% points, the blue line represents the mean quality. **Figure S14.** Kinship relationships among the 94 *P. tremula* individuals. The values of the relatedness statistics were calculated according to the method implemented in GEMMA. **Figure S15.** Two-dimensional distribution for squared loadings *ρ*^2^_j1_ with the first environmental principal component estimated from PCAdapt and Weir & Cockerham’s *F*_ST_ values of the common SNPs with minor allele frequency higher than 5%. The yellow to dark blue to light blue gradient indicates decreased density of observed events at a given location in the graph. Black dots represent SNPs fulfilling the significance threshold requirement defined by PCAdapt. **Figure S16.** (a) Correlations between pairs of the 39 environmental variables. Blue indicates a positive relationship, and red indicates a negative relationship. Color intensity is proportion to Pearson’s correlation coefficient. (b) The percent of explained variance for each principal component (PC) from the principal component analysis (PCA) of all 39 environmental variables. (c) Biplot for all environmental variables loaded on the top two PCs. (d) The relationship between scores of the first environmental principal component (PC) and the length of growing season, which is represented as the number of days with temperature higher than 5 °C, for samples in the 12 populations of *P. tremula*. **Figure S17.** Comparison between imputation accuracy with minor allele frequency (MAF) under a simulation test. Color lines shows imputation accuracy compared with MAF under various ratio of artificial missed SNPs had been imputed by BEAGLE v 4.1. **Figure S18.** Distribution of maximum distance between significant composite likelihood ratio (CLR) peaks calculated using the simulated data from SweepFinder2. The dotted line denotes the 95% quartile and the dark red line indicates the value calculated from the empirical dataset in this study. **Figure S19.** Key used for scoring bud set in the field experiment with transgenic *PtFT2* lines at Våxtorp, Sweden

### Additional file4

**Table S3.** List of the 1615 candidate SNPs associated with local adaptation.

### Additional file5

**Table S4.** Statistic summary (median and central 95% range) for five selective sweep measures across the ~700 kb region around *PtFT2* gene on chromosome 10 and genome-wide level. Pairwise nucleotide diversity (π), genetic divergence between groups of populations (*F*_ST_), H12, H2/H1, and composite likelihood ratio (CLR) test are compared for three groups of populations, South (pop 1-6), Mid (pop 7-8) and North (pop 9-12) corresponding to Fig. 4.

### Additional file6

**Table S5.** Anova tables for analyses of gene expression in greenhouse and common garden experiments

### Additional file7

**Table S6.** Average values of 39 environmental variables over the years 2002-2012 for the original sample location of 94 *Populus tremula* individuals used in this study.

